# Voltage to Calcium Transformation Enhances Direction Selectivity in *Drosophila* T4 neurons

**DOI:** 10.1101/2022.07.01.498438

**Authors:** Abhishek Mishra, Alexander Borst, Juergen Haag

## Abstract

A critical step in neural information processing is the transformation of membrane voltage into calcium signals leading to transmitter release. However, the effect of voltage to calcium transformation on neural responses to different sensory stimuli is not well understood. Here, we use in vivo two-photon imaging of genetically encoded voltage and calcium indicators, Arclight and GCaMP6f respectively, to measure responses in *Drosophila* direction-selective T4 neurons.

Comparison between Arclight and GCaMP6f signals revealed calcium signals to have a significantly higher direction selectivity compared to voltage signals. Using these recordings we build a model which transforms T4 voltage responses to calcium responses. The model reproduces experimentally measured calcium responses across different visual stimuli using different temporal filtering steps and a stationary non-linearity. These findings provide a mechanistic underpinning of the voltage-to-calcium transformation and show how this processing step, in addition to synaptic mechanisms on the dendrites of T4 cells, enhances direction selectivity in the output signal of T4 neurons.

## Introduction

In order to guide animal behavior, neurons perform a wide range of computations. Neurons encode information via graded changes in membrane potential or action potential frequency. Mostly they communicate via chemical synapses which requires the release of neurotransmitters. When the presynaptic membrane is sufficiently depolarized, voltage-gated calcium channels open and allow *Ca*^2+^ to enter the cell (Luo 2020). Calcium entry leads to the fusion of synaptic vesicles with the membrane and release of neurotransmitter molecules into the synaptic cleft (Chapman 2002). As neurotransmitters diffuse across the synaptic cleft, they bind to receptors in the postsynaptic membrane, causing postsynaptic neuron to depolarize or hyperpolarize, passing the information from pre to postsynaptic neurons (Di Maio 2008). Voltage to calcium transformation therefore represents one crucial step in neural information processing and neural computation.

A classic example of neural computation is how *Drosophila* neurons compute the direction of visual motion (Borst *et al*. 2020). In *Drosophila*, visual information is processed in parallel ON (contrast increments) and OFF (contrast decrements) pathways (Joesch *et al*. 2010; Eichner *et al*. 2011). Direction selectivity emerges three synapses downstream of photoreceptors, in T4 and T5 for ON and OFF pathways respectively. Four subtypes of T4 and T5 cells exist, each responding selectively to one of the four cardinal directions (Maisak *et al*. 2013). Amazingly, right at the first stage where direction selectivity emerges, T4 and T5 cells exhibit a high degree of direction selectivity, with no responses to null direction stimuli. This statement is, however, based on calcium recordings. Whole-cell patch clamp recordings show a somewhat different picture: While preferred direction stimuli also lead to large membrane depolarizations, edges or gratings moving along the null directions elicit smaller but significant responses as well (Wienecke *et al*. 2018; Groschner *et al*. 2022). This hints to an additional processing step where voltage signals are transformed into calcium signals that increases direction selectivity of the cells. In order to study this step systematically, we recorded both voltage and calcium signals in response to a large stimulus set that includes gratings and edges moving along various directions at different speeds and contrasts. Using these data, we build a model that captures the transformation from voltage to calcium by a few linear and non-linear processing steps.

## Results

We first expressed the genetically encoded calcium indicator GCaMP6f (Chen *et al*. 2013) in T4 cells projecting to layer 3 of the lobula plate. These cells have upward motion as their preferred direction (PD) and downward motion as their null direction (ND). We also expressed the genetically encoded voltage indicator Arclight (Jin *et al*. 2012) using the same driver line. Arclight’s fluorescence decreases with membrane depolarization and increases with membrane hyperpolarization. To compare the voltage and calcium signals, we recorded the neural activity in T4c cells dendrites in medulla layer 10 in response to the same set of stimuli using 2-photon microscopy (Denk *et al*. 1990). The complete stimuli set included square-wave gratings of 30° spatial wavelength moving in 12 different directions, and ON edges moving in PD and ND, at four different speeds (15°*s*^−1^, 30°*s*^−1^, 60°*s*^−1^, 120°*s*^−1^) and four different contrasts (10%, 20%, 50%, 100%).

In a first set of experiments, we measured the voltage and calcium signals in response to gratings moving in PD and ND at four different speeds (figure 1A). As the grating stimuli consists of alternate bright and dark bars moving in a certain direction, there was a modulation in the Arclight (black traces) and GCaMP6f (red traces) responses to it. The GCaMP6f responses showed modulations only for slower speeds, while Arclight responses revealed modulations also for faster speeds. The response amplitudes were much higher for GCaMP6f (≈ 2.0Δ*F*/*F*) compared to Arclight (≈ −0.06Δ*F*/*F*). The peak responses (maximum Δ*F*/*F*) decreased with increasing stimulus speed both for GCaMP6f and Arclight (figure 1B). To understand if voltage to calcium transformation affects direction selectivity in T4 cells, we compared the responses to gratings moving in PD and ND. GCaMP6f responses in ND were negligible compared to its responses in PD, while for Arclight responses in ND were quite visible. We quantified the direction selectivity using a direction selectivity index (DSI) calculated as the difference of the peak responses to preferred and null direction, divided by the sum of the peak responses (Materials and Methods equation (1)). The results revealed a high degree of direction selectivity of ≈ 0.8 for GCaMP6f at slower velocities, compared to a direction selectivity of ≈ 0.4 for Arclight (figure 1E). For both GCaMP6f and Arclight signals, direction selectivity decreased with increasing velocity.

**Figure 1.**
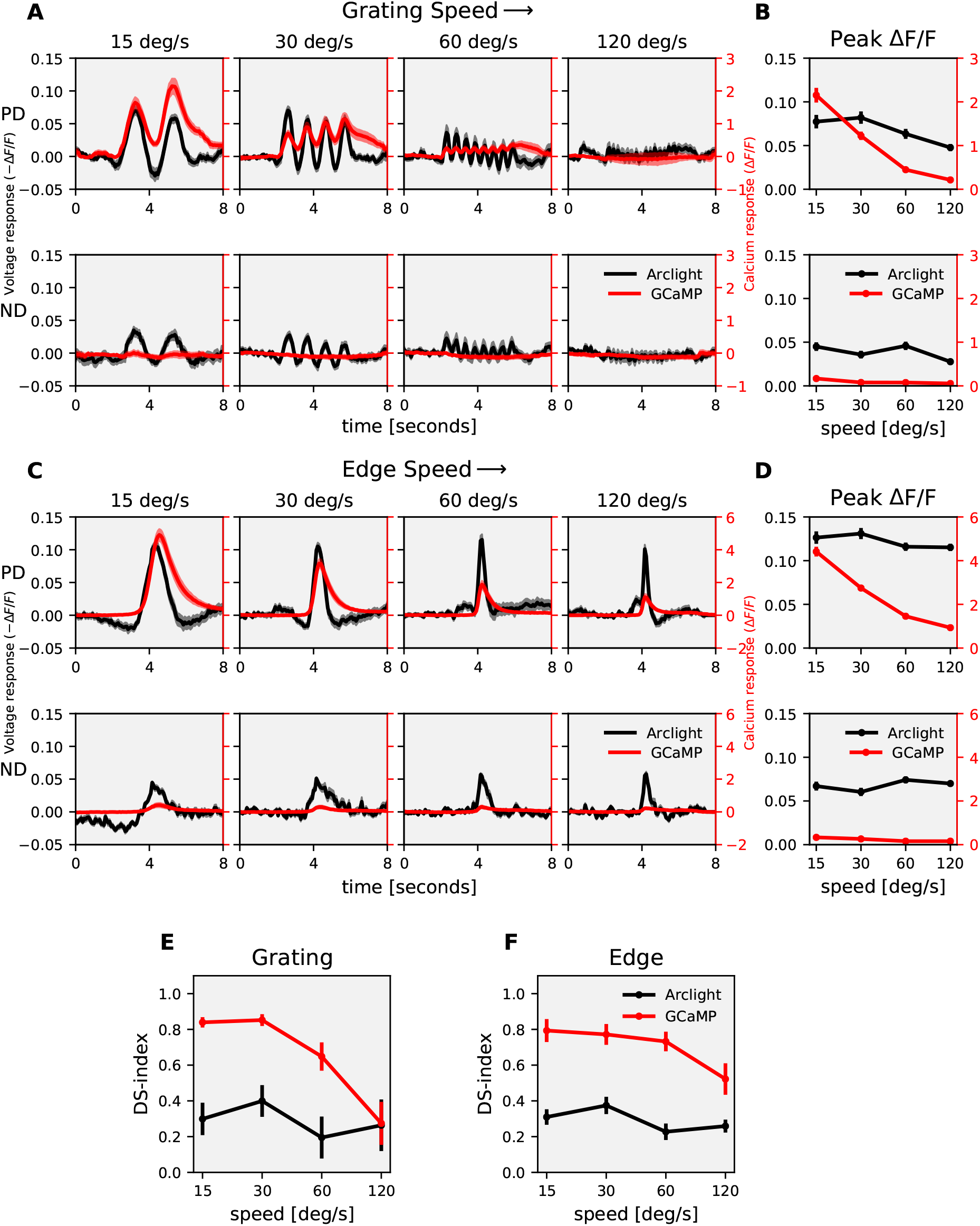
T4c speed dependence : (A) T4c Arclight (black) and GCaMP6f (red) responses to grating moving in PD (top row) and ND (bottom row) at 4 different speeds. The plots have twin y-axis. The left y-axis of the plot represents voltage responses i.e. changes in Arclight fluorescence (−Δ*F*/*F*) and the right y-axis of the plot represents calcium responses i.e. changes in GCaMP6f fluorescence (Δ*F* /*F*) (B) T4c peak responses to grating moving in PD (top) and ND (bottom) at 4 different speeds. (n = 20 ROIs from N = 10 flies for Arclight, n = 18, N = 9 for GCaMP6f) (C) T4c Arclight (black) and GCaMP6f (red) responses to ON-edge moving in PD (top row) and ND (bottom row) at 4 different speeds. (D) T4c peak responses to ON-edge moving in PD and ND at 4 different speeds. (n = 29, N = 10 for Arclight, n = 17, N = 4 for GCaMP6f) (E) Direction selectivity index (DSI) calculated as difference of peak responses in PD and ND divided by the sum of peak responses for grating. (F) Direction selectivity index (DSI) for ON-edge. All data shows the mean ± SEM. PD: preferred direction, ND: null direction.

Next, instead of gratings, we used moving bright edges with all other stimulus parameters remaining the same (figure 1C). As the edge moves upward on the screen, it crosses the receptive field of T4c neurons (≈ 15°) only once. Hence, there was only a single peak in the response. The peak response decreased with increasing stimulus speed for GCaMP6f, while the peak response remained almost constant for Arclight throughout all speeds (figure 1D). When comparing edge responses moving along preferred and null directions, GCaMP6f showed negligible responses in null direction while Arclight revealed considerable responses to null direction stimuli. The direction selectivity index was again much higher for GCaMP6f compared to Arclight (figure 1F). Together these results show that GCaMP6f signals have a high level of direction selectivity compared to Arclight signals, both for grating and edge stimuli.

The stimulus strength was further varied by changing the contrast between bright and dark bars for gratings and between moving edge and background for edge stimuli. We measured Arclight and GCaMP6f responses to gratings moving at 30°*s*^−1^ at four different contrasts (figure 2A). Increasing contrast resulted in an increase in response for both Arclight and GCaMP6f. GCaMP6f signals were modulated at the temporal frequency of the grating but showed an additional rise over time. This slow increase was not observed in Arclight signals. We also measured Arclight and GCaMP6f responses to ON edges moving at the same speed of 30°*s*^−1^ but having different contrasts (figure 2C). The peak response (maximum Δ*F*/*F*) increased with increasing contrast (figure 2D). Similar to previous experiments, the direction selectivity index was much higher for GCaMP6f (≈ 0.9) compared to that for Arclight (≈ 0.4) (figure 2E,F).

**Figure 2.**
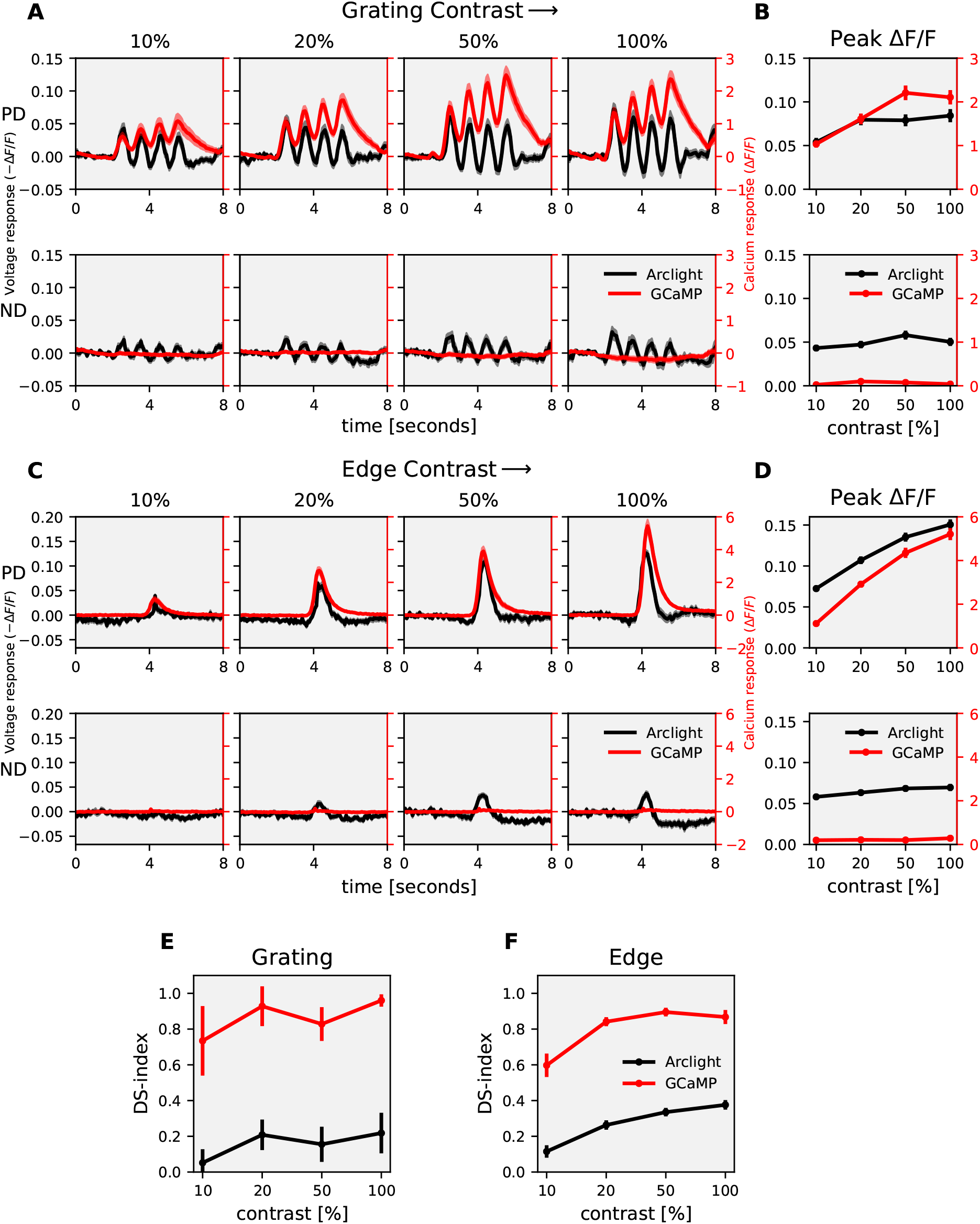
T4c contrast dependence : (A) T4c Arclight (black) and GCaMP6f (red) responses to grating moving in PD (top row) and ND (bottom row) at 4 different contrasts. The left y-axis of the plot represents voltage responses i.e. changes in Arclight fluorescence (−Δ*F*/*F*) and the right y-axis of the plot represents calcium responses i.e. changes in GCaMP6f fluorescence (Δ*F*/*F*) (B) T4c peak responses to grating moving in PD (top) and ND (bottom) at 4 different contrasts. (n = 23 ROIs from N = 11 flies for Arclight, n = 22, N = 9 for GCaMP6f) (C) T4c Arclight (black) and GCaMP6f (red) responses to ON-edge moving in PD (top row) and ND (bottom row) at 4 different contrasts. (D) T4c peak responses to ON-edge moving in PD and ND at 4 different contrasts. (n = 36, N = 5 for Arclight, n = 41, N = 7 for GCaMP6f) (E) Direction selectivity index (DSI) calculated as difference of peak responses in PD and ND divided by the sum of peak responses for grating. (F) Direction Selectivity Index (DSI) for ON-edge. All data shows the mean ± SEM.

In the results presented so far we compared responses for two directions only, i.e. along the preferred (upward) and along the null direction (downward). We next extended the comparison to motion along 12 directions, from 0° to 360° in steps of 30°. For this comparison, we determined the normalized peak responses of Arclight and GCaMP6f signals to gratings moving in 12 directions at 4 different speeds and 4 different contrasts, respectively (figure 3A, B). The directional tuning was much sharper for GCaMP6f compared to Arclight. To quantify this we calculated the directional tuning index *L*_*dir*_ (Mazurek *et al*. 2014) for each speed and each contrast as the vector sum of the peak responses divided by the sum of all individual vector magnitudes (Materials and Methods equation (2)). In general, the directional tuning indices again were much higher for GCaMP6f (≈ 0.6) compared to that of Arclight (≈ 0.2) (figure 3C, D). Together these results show that GCaMP6f signals have a higher degree of directional tuning across different speeds and contrasts than Arclight.

**Figure 3.**
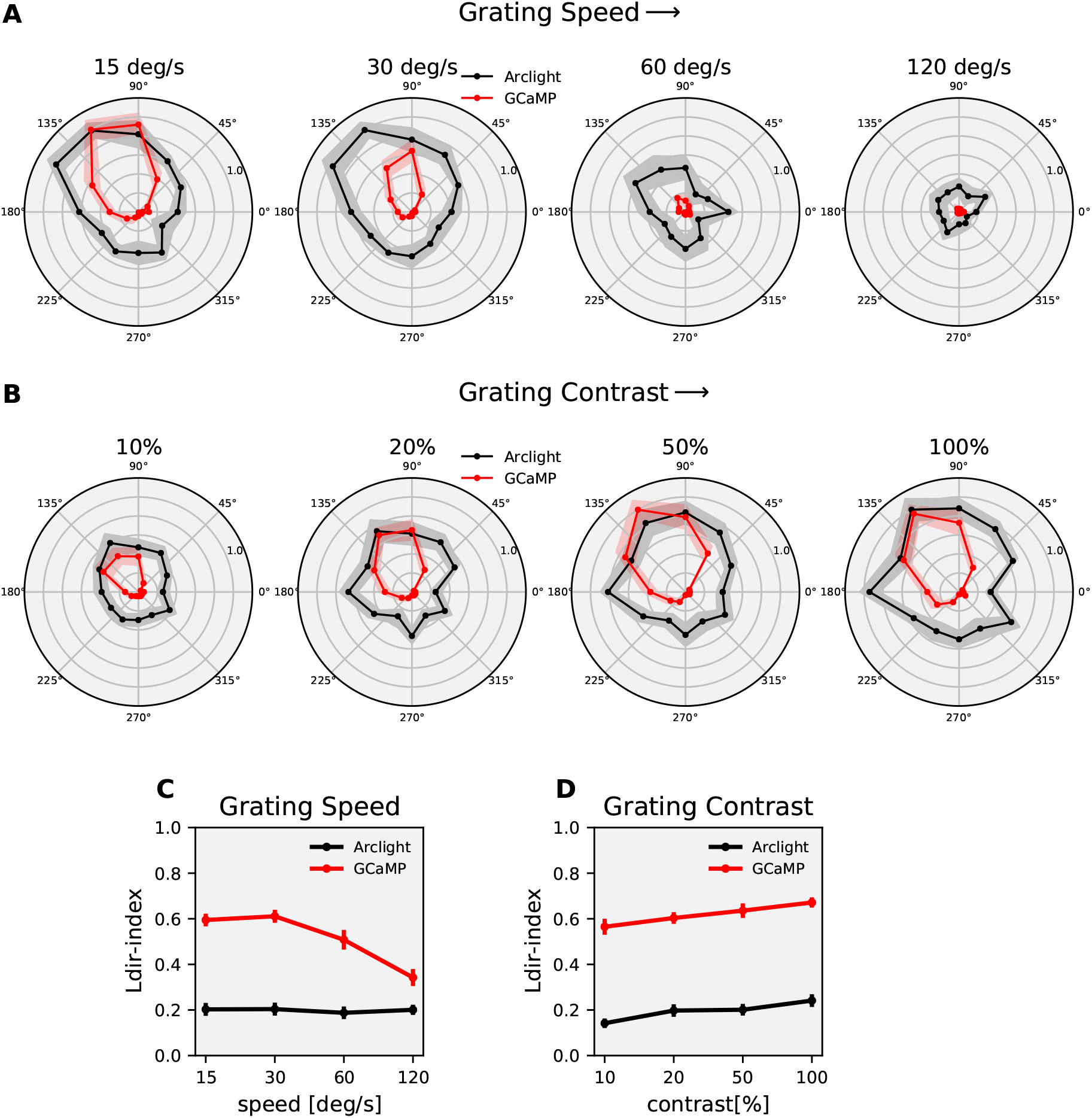
T4c direction tuning : (A) T4c Arclight (black) and GCaMP6f (red) normalized peak responses to grating moving in 12 directions at 4 different speeds. (n = 20 ROIs from N = 10 flies for Arclight, n = 18, N = 9 for GCaMP6f) (B) T4c Arclight (black) and GCaMP6f (red) normalized peak responses to grating moving in 12 directions at 4 different contrasts. (n = 23, N = 11 for Arclight, n = 22, N = 9 for GCaMP6f) (C) The directional tuning index *L*_*dir*_ for grating moving at 4 different speeds. The directional tuning index is calculated as the vector sum of the peak responses divided by the sum of all individual vector magnitudes. (D) The directional tuning index for grating at 4 different contrasts. All data shows the mean ± SEM measured in 5 different flies.

How does the voltage to calcium transformation lead to calcium signals with significantly higher directional tuning compared to voltage signals? To address this question, we constructed an algorithmic model (figure 4) which takes Arclight signals as inputs and outputs GCaMP signal. In order to find the optimal parameter values, we first defined an error function. For each stimulus condition, the error was calculated as the sum of the squared difference between the model and experimental data at each time-point (Materials and Methods equation (3)). There were a total of 112 stimulus conditions: gratings speed (48), gratings contrast (48), edge speed (8) and edge contrast (8). The total error amounted to the sum of errors across all stimulus conditions (Materials and Methods equation (4)). We defined the model error as the total error divided by the power of the data (Materials and Methods equation (5)). We then found the optimal parameters values of the model that correspond to the minimum total error using Python SciPy optimize minimize function (Virtanen *et al*. 2020).

**Figure 4.**
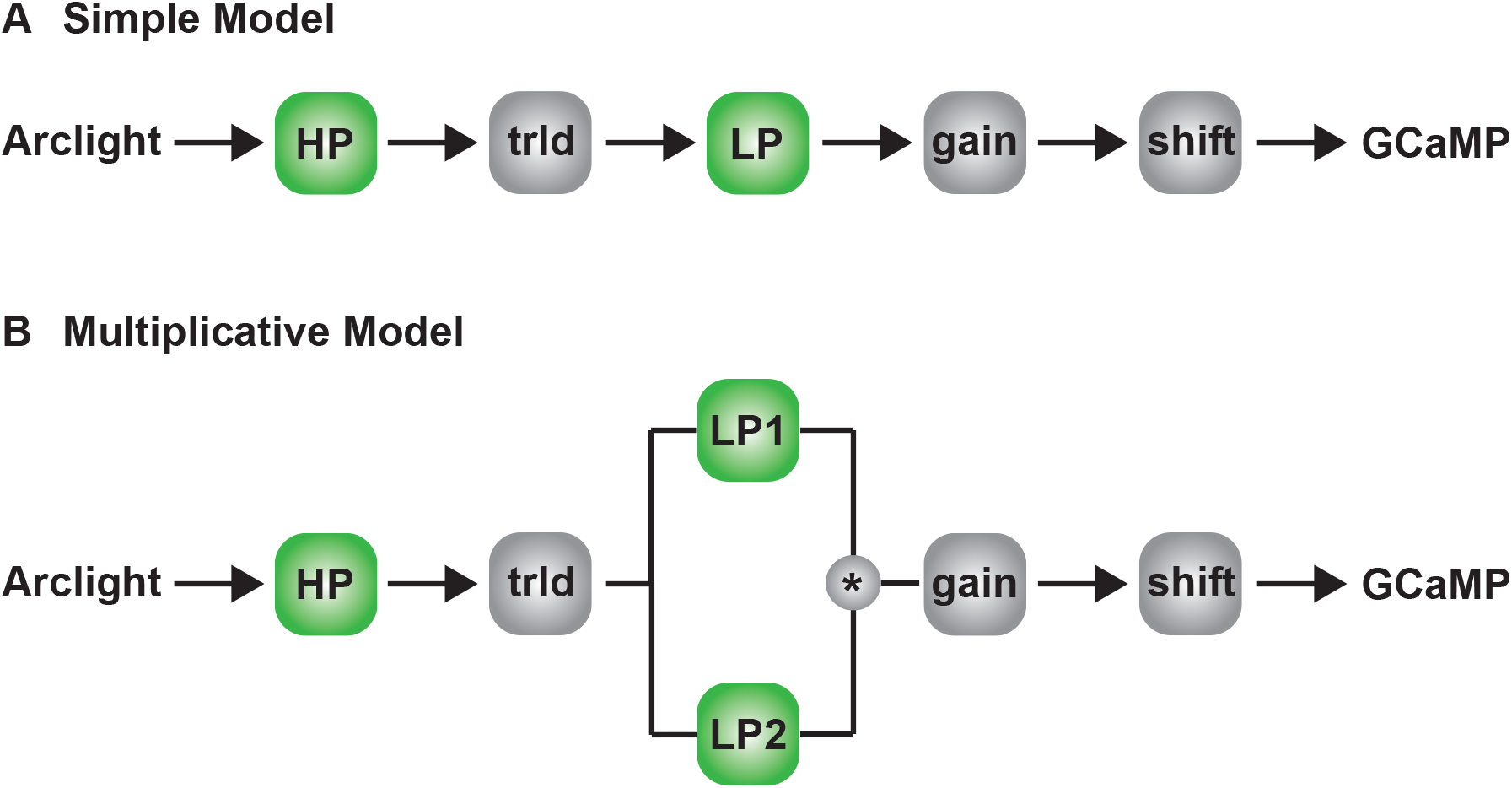
Models for voltage to calcium transformation : (A) Simple model consisting of High-Pass filter (HP), threshold (trld), Low-Pass filter (LP), gain and shift. (B) Multiplicative model combining output of two low-pass filters via multiplication.

We started with a simple model (figure 4A). The model first passes the Arclight signal through a high-pass filter. The high-pass filter brings the input Arclight signal closer to the actual voltage signal by removing the slowly fluctuating Arclight indicator dynamics. This is followed by a threshold, assuming that the voltage changes below a certain threshold does not affect the calcium level in the cell. Now, few experimental observations which we took into consideration for building up the model further were as follows : First, the GCaMP6f response to gratings showed modulations only for slower speeds, whereas Arclight response had modulations even at faster speeds (figure 1A). This suggests that the GCaMP6f signal is a low-pass filtered version of the Arclight signal. In the simple model, we used a single low-pass filter followed by a gain and time-shift. Multiplication with a gain factor was required since GCaMP6f signals have a much higher magnitude compared to Arclight. Arclight and GCaMP6f responses were recorded from cells in different flies with different receptive fields, therefore the responses had different phases, and a time-shift was necessary to align the signals. However, the simple model with single low-pass filter could not reproduce responses across all stimuli. The model error for the complete dataset fit for the simple model was around 34%. Specifically, the simple model failed to suppress the ND-responses and to reproduce the edge responses. The directional tuning index *L*_*dir*_ was much smaller for the simple model compared to the experimental data (figure 5E,F). Second, the GCaMP6f responses in addition to modulation also had a steady rise over time whereas Arclight signal only had modulations (figure 1A, 2A). For reproducing the edge responses and modulation in grating responses, the model needed a low-pass filter with a small time constant. However to simulate the steady rise in the grating signal, a low-pass filter with a large time constant was necessary. Hence, we combined the output of two low-pass filters. Summing up the low-pass filter outputs did not lead to much improvement.

**Figure 5.**
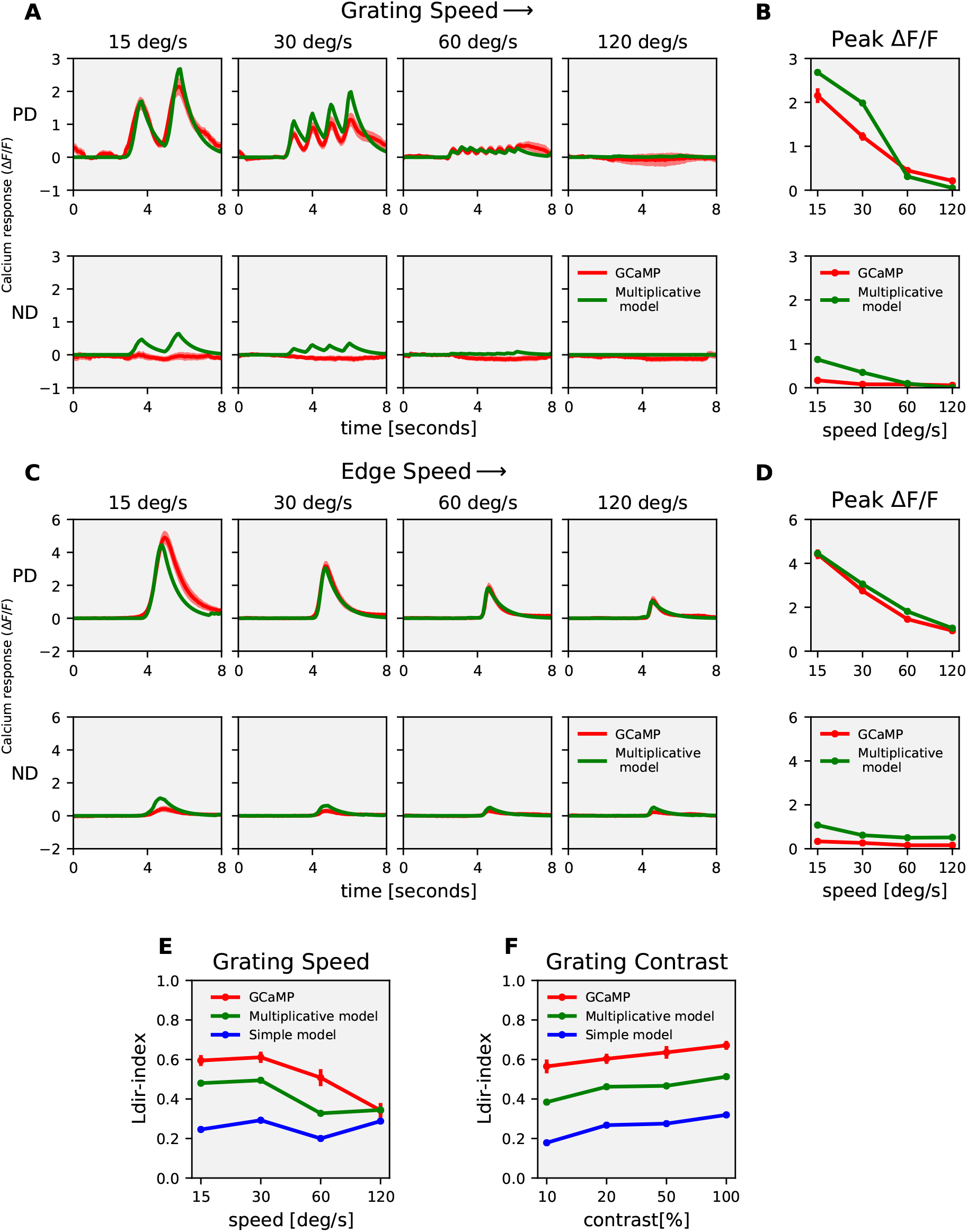
Model responses : (A) T4c GCaMP6f (red) and multiplicative model (green) responses to grating moving in PD (top row) and ND (bottom row) at 4 different speeds. (B) T4c GCaMP6f and model peak responses to grating moving in PD (top) and ND(bottom) at 4 different speeds. (C) T4c GCaMP6f (red) and multiplicative model (green) responses to ON-edge moving in PD (top row) and ND (bottom row) at 4 different speeds. (D) T4c GCaMP6f and model peak responses to ON-edge moving in PD (top) and ND (bottom) at 4 different speeds. (E, F) The directional tuning index *L*_*dir*_ for GCaMP6f (red), multiplicative (green) and simple (blue) model for grating moving in 12 directions at 4 different speeds and at 4 different contrasts respectively.

However, combining both outputs from the low-pass filters with a multiplication led to significant decrease in the error. The model error for the multiplicative model (figure 4B) then was only at around 20%.

The multiplicative model thus has in total 6 parameters - high-pass filter time constant, threshold, low-pass filter 1 time constant, low-pass filter 2 time constant, gain and shift. The multiplicative model was able to reproduce calcium signals across different visual stimuli (figure 5). It could reproduce both the modulation as well as slow rise in the GCaMP6f signal in response to gratings (figure 5A). The multiplicative model could also reproduce the ON edge speed tuning responses across different speeds (figure 5C,D). The directional tuning index *L*_*dir*_ were similar for multiplicative model and experimental data across slower speeds and all contrasts (figure 5E,F).

Is the slow rise in GCaMP6f signals over time due to the properties of T4 cells or due to the properties of GCaMP6f? To answer this question we used a faster version of the calcium indicator GCaMP8f (Zhang *et al*. 2020). GCaMP8f was expressed in T4c cells using the same driver line. The experiments were repeated using grating stimuli in 12 directions at 4 speeds and ON edges moving in PD and ND. T4c cells GCaMP8f responses were similar to GCaMP6f responses but faster. As with GCaMP6f, GCaMP8f signals had modulation and slow rise over time. We further compared the model parameters values for GCaMP6f data fit and GCaMP8f data fit (figure 6). The model parameters were similar, but with time constants having smaller values for GCAMP8f as it is a faster indicator. Therefore, the slow rise in the calcium signal is not due to the properties of GCaMP6f indicator.

**Figure 6.**
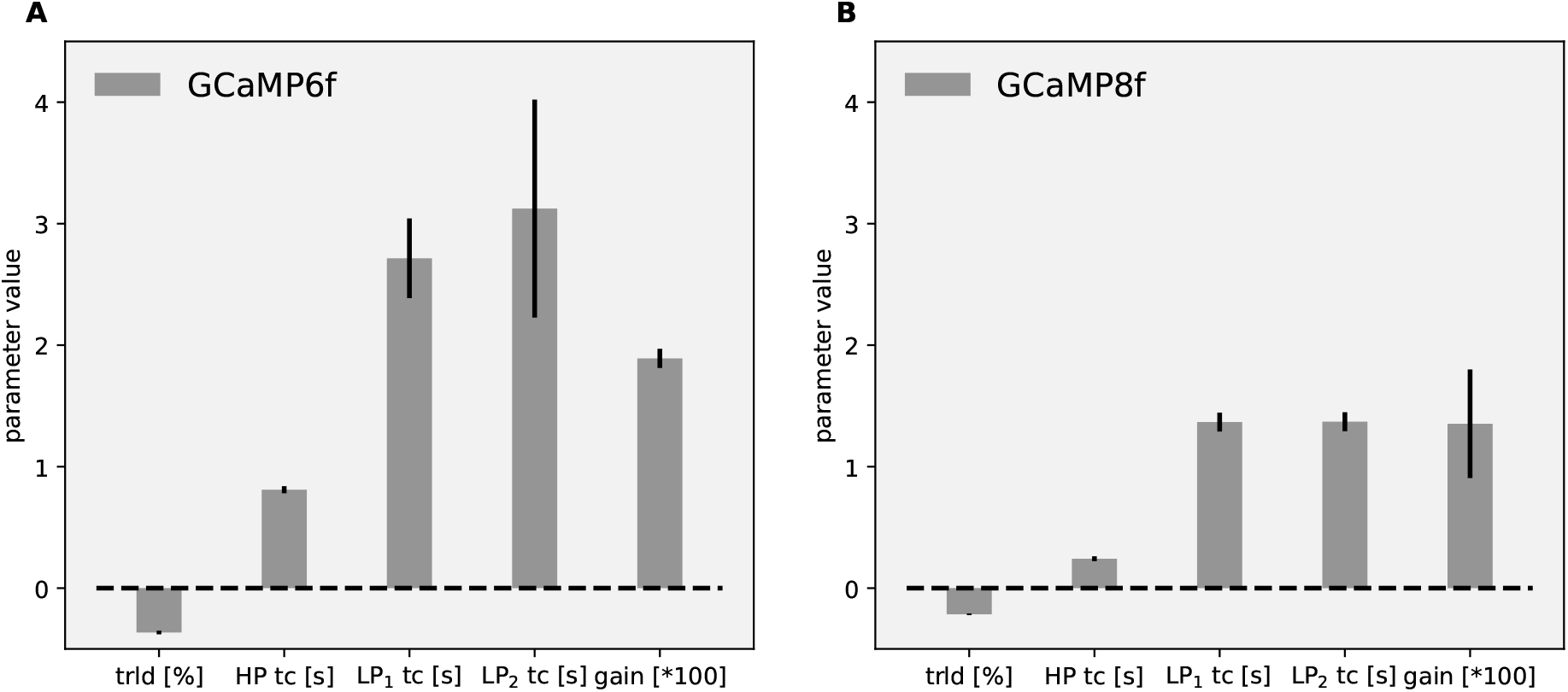
Model parameters for GCaMP6f (A) and GCaMP8f (B) : Data shows mean ± SD for optimal parameters for the multiplicative model. The data were fit for grating moving in 12 directions and 4 speeds, and for ON-edge moving in PD and ND at 4 speeds. trld : threshold, HP : High Pass, LP : Low Pass, tc : Time constant

To reproduce the calcium responses for direction-selective T4c cells under all stimuli conditions, a multiplicative model was required. For non-direction-selective cells, what does the voltage to calcium transformation look like, and is the simple model able to replicate the calcium response for these cells? In order to answer this question, we expressed Arclight & GCaMP6f in medulla neurons Mi1 & Tm3 cells, which are both non-direction-selective. Mi1 and Tm3 are pre-synaptic to T4 cells and have ON-center receptive field (Behnia *et al*. 2014; Arenz *et al*. 2017). We measured Mi1, Tm3 Arclight (black) and GCaMP6f (red) responses to gratings moving at 4 different speeds and to gratings moving at 4 different contrasts (figure 7). The gratings were moved in only one direction, since the direction does not affect non-direction-selective cells’ responses. Contrary to T4, Mi1 GCaMP6f responses had only modulation without a slow increase over time (figure 7A). Tm3 GCaMP responses did not increase over time, and showed only modulation for gratings moving at 15°*s*^−1^. For gratings moving at 30°*s*^−1^ and 60°*s*^−1^, there was an increase in Tm3 GCaMP6f response over time, but the Arclight response also already had a slow increment over time (figure 7A). Similar to T4, the peak response for Mi1 and Tm3 decreased with an increase in speed and increased with an increase in contrast (figure 7B, D). Together, these results show that voltage to calcium transformation causes GCaMP6f response increment over time only for direction-selective T4 cells and not for non-direction-selective Mi1 and Tm3 cells.

**Figure 7.**
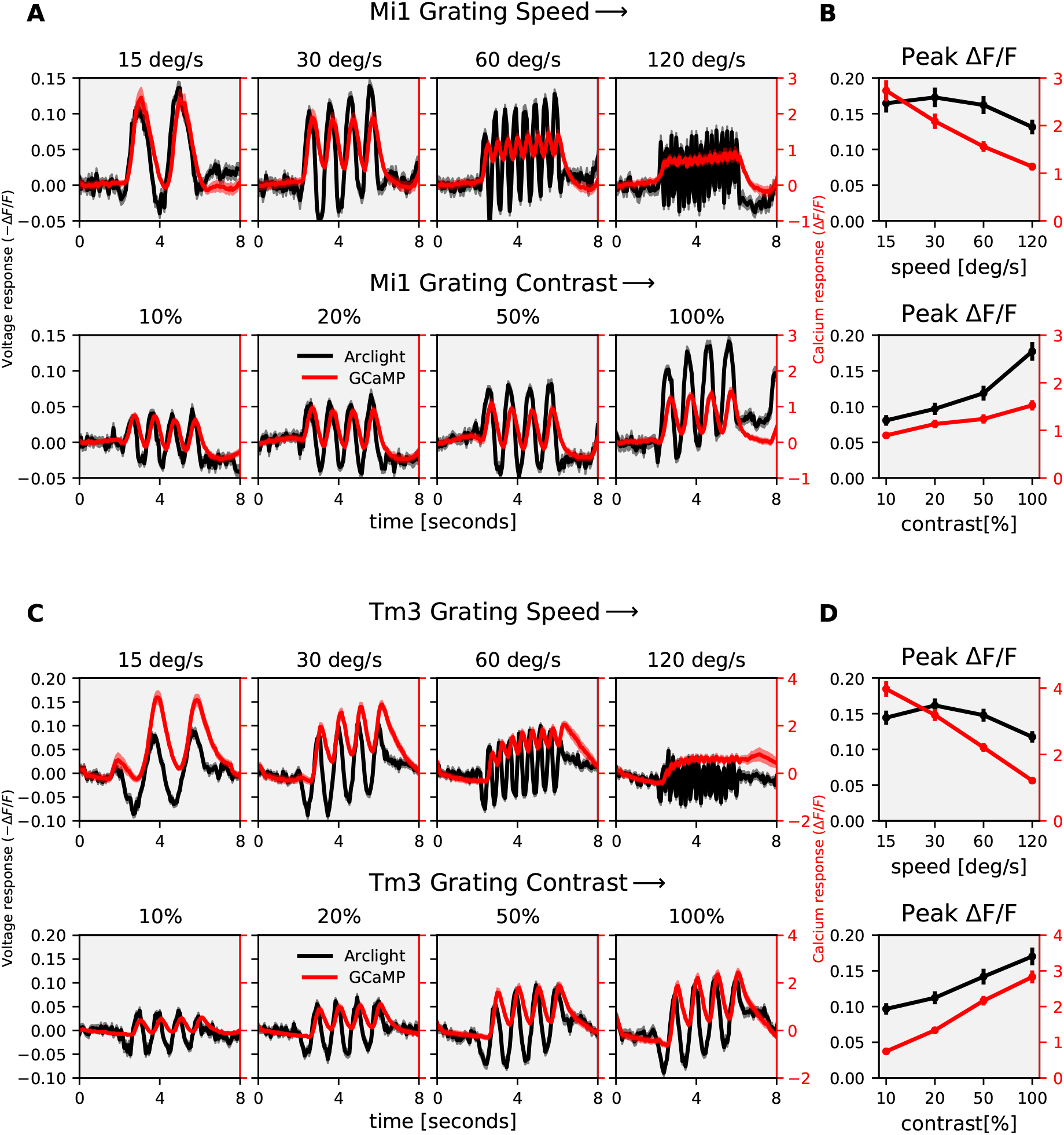
Mi1, Tm3 speed and contrast dependence : (A) Mi1 Arclight (black) and GCaMP6f (red) responses to grating moving at 4 different speeds (top row) and 4 different contrasts (bottom row). The left y-axis of the plot represents voltage responses i.e. changes in Arclight fluorescence (−Δ*F*/*F*) and the right y-axis of the plot represents calcium responses i.e. changes in GCaMP6f fluorescence (Δ*F*/*F*) (B) Mi1 peak responses to grating moving at 4 different speeds (n = 24 ROIs from N = 5 flies for Arclight, n = 19, N = 8 for GCaMP) and 4 different contrasts (n = 24, N = 5 for Arclight, n = 22, N = 8 for GCaMP). (C) Tm3 Arclight (black) and GCaMP6f (red) responses to grating moving at 4 different speeds (top row) and 4 different contrasts (bottom row). (D) Tm3 peak responses to grating moving at 4 different speeds (n = 52, N = 5 for Arclight, n = 37, N = 4 for GCaMP) and 4 different contrasts (n = 35, N = 5 for Arclight, n = 36, N = 4 for GCaMP). All data shows the mean ± SEM.

Next, we used the model described in figure 4 to reproduce Mi1 and Tm3 calcium responses using their Arclight responses. As discussed earlier, the simple model (figure 4A) with single low-pass filter was not able to reproduce T4 calcium responses across all stimuli. However, for Mi1 and Tm3, the simple model with a single low-pass filter was able to reproduce the calcium responses across all stimuli conditions (figure 8). The model also accurately replicated the speed and contrast tuning for Mi1 and Tm3 (figure 8B, D). We further compared the model error for simple and multiplicative model for Mi1, Tm3 and T4c data (figure 9). The model error for Mi1 and Tm3 for simple model was ≈ 6.5% and ≈ 5.9% respectively compared to ≈ 11.9% and ≈ 7% for the multiplicative model. Thus, the simple model already performed well for Mi1 and Tm3 dataset, and changing to multipicative model did not improve the performance. For the T4c dataset the model error was ≈ 34% and ≈ 21% for the simple and multiplicative model respectively. Hence, the multiplicative model with two low-pass filters performed better for T4c dataset whereas for Mi1 and Tm3 the Simple model with single low-pass filter was sufficient to reproduce the calcium responses. This suggests that voltage-to-calcium transformation is more complex for direction-selective cell T4 than for the non-direction-selective cells Mi1 and Tm3.

**Figure 8.**
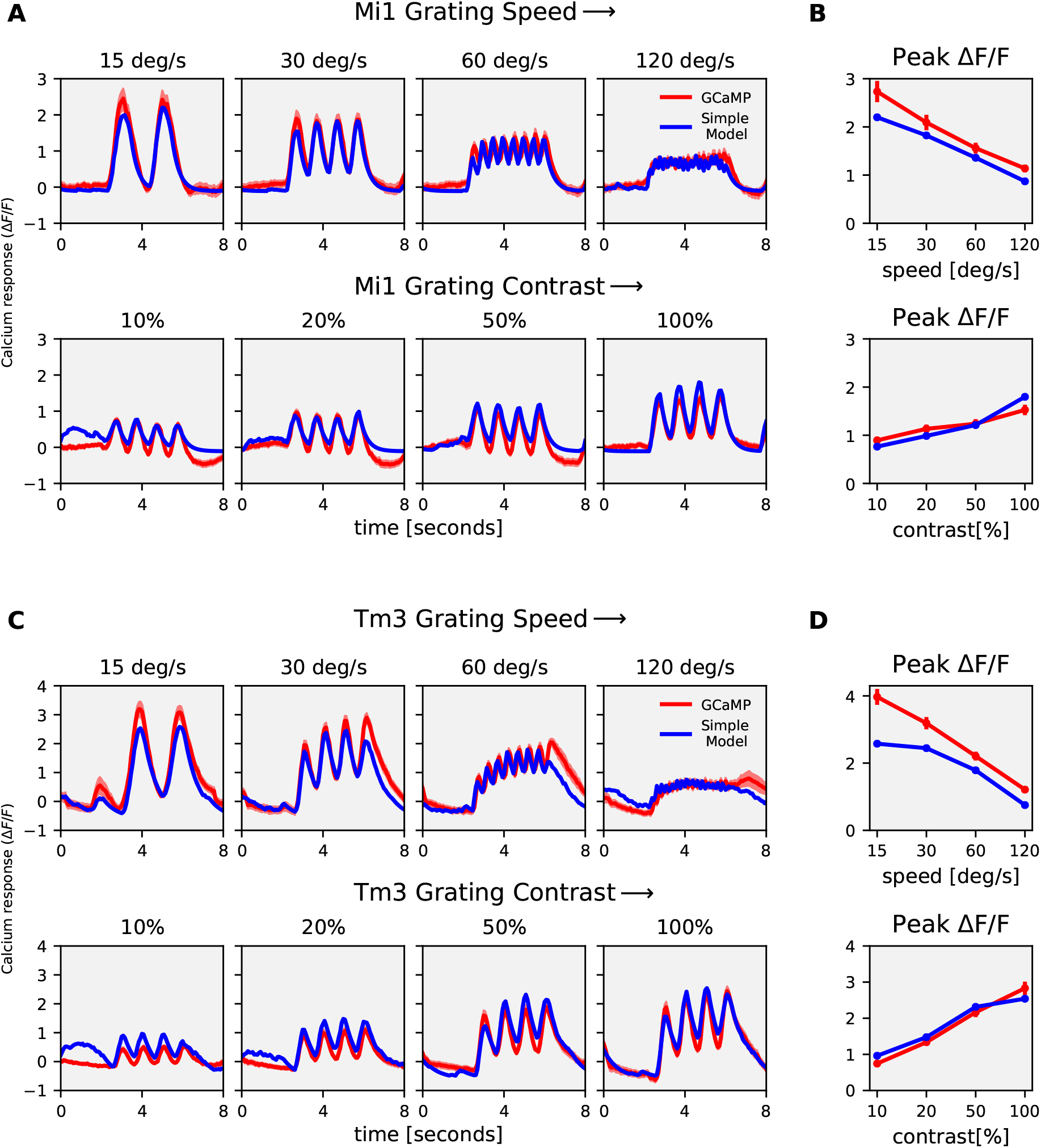
Mi1, Tm3 Simple model responses : (A) Mi1 GCaMP6f (red) and simple model (blue) responses to gratings moving at 4 different speeds (top row) and to gratings moving at 4 different contrasts (bottom row). (B) Mi1 GCaMP6f and model peak responses to gratings moving at 4 different speeds (top) and 4 different contrasts (bottom). (C) Tm3 GCaMP6f (red) and simple model (blue) responses to gratings moving at 4 different speeds (top row) and to gratings moving at 4 different contrasts (bottom row). (D) Tm3 GCaMP6f and model peak responses to gratings moving at 4 different speeds (top) and 4 different contrasts (bottom).

**Figure 9.**
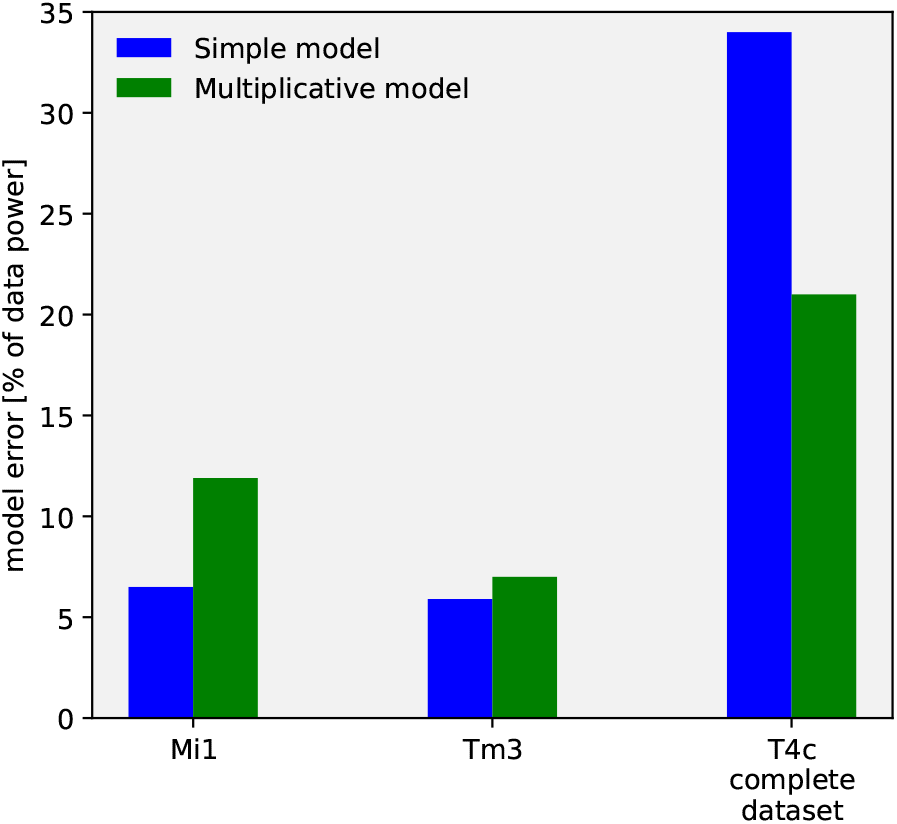
Model error for the simple and multiplicative model : The model error for the simple model (blue) and multiplicative model (green). Mi1 and Tm3 dataset consists of gratings at 4 different speeds and contrast moving in a single direction. T4c complete dataset consists of gratings moving in 12 different directions, and ON edge moving in PD, ND at 4 different speeds and contrasts i.e. a total of 112 stimuli conditions.

## Discussion

Neuronal signaling and information processing involves the transformation of membrane voltage into calcium signals, which lead to transmitter release. Computations can occur at different stages in the signaling cascade: 1.) dendritic integration and processing of voltage signals. 2.) transformation from voltage to calcium and 3.) between calcium and neurotransmitter release. In this study, we explored the transformation of voltage to calcium in T4-cells, the first direction-selective neurons in the *Drosophila* ON motion vision pathway.

We found that the voltage to calcium transformation in T4c neurons enhances their direction selectivity: calcium signals in T4c cells have a significantly higher direction selectivity and tuning compared to membrane voltage across different stimuli conditions (figure 1-3). The direction selectivity index for calcium signals compared with voltage signals for a few stimuli conditions was previously found to be higher in a study in T5 cells using ASAP2f as an optical voltage indicator (Wienecke *et al*. 2018). As calcium is required for neurotransmitter release, this is expected to increase the direction selectivity of T4/T5 cells’ output signals. In the lobula plate, T4/T5 cells provide inputs onto large lobula plate tangential cells that are depolarized during preferred and hyperpolarized during null direction motion (Mauss *et al*. 2014). For example, vertical system (VS) cells with dendrites in layer 4 receive direct excitatory inputs from downward tuned T4d/T5d neurons causing depolarization during motion in the downward preferred direction. These VS cells also receive indirect inhibitory inputs from upward tuned T4c/T5c neurons via glutamatergic LPi3-4 neurons projecting from layer 3 to layer 4 causing hyperpolarization in VS cells during motion in the upward null direction. Upon silencing LPi3-4 neurons’ synaptic output via tetanus toxin, VS neurons depolarization response in the preferred direction did not change, but the null direction response was absent (Mauss *et al*. 2015). This suggests T4/T5 do not release any transmitter in response to null direction motion, which matches our findings for the calcium responses. Thus, voltage to calcium transformation increases direction selectivity in T4/T5 cells and this enhances direction selectivity in downstream neurons.

Electrophysiology has been the most frequently used method to measure the membrane potential changes in neurons. However, due to the small size of neurons in the optic lobe, single-cell electrophysiological recordings of these neurons have been difficult. Genetically encoded voltage indicators (GEVIs) have evolved as powerful tools for recording changes in neuronal membrane potentials (Yang *et al*. 2016). Optical methods of monitoring brain activity are appealing because they allow simultaneous, noninvasive monitoring of activity in many individual neurons. We used a fluorescence protein (FP) voltage sensor called Arclight (Jin *et al*. 2012). Arclight is based on the fusion of the voltage-sensing domain of *Ciona intestinalis* voltage-sensitive phosphatase (Murata *et al*. 2005) and the fluorescent protein super ecliptic pHluorin with an A227D mutation. Arclight has been shown to robustly report both subthreshold events and action potentials in genetically targeted neurons in the intact *Drosophila* brain (Cao *et al*. 2013).

We built a model to capture voltage to calcium transformation in T4c, Mi1, and Tm3 cells. A simple model with a single low-pass filter was able to reproduce calcium responses in non-direction-selective Mi1 and Tm3 cells (figure 8), whereas a more complex model combining the output of two low-pass filters via a multiplication was required to reproduce T4c calcium responses (figure 5). The direction selectivity for the simple model signals for T4c was lower compared to the multiplicative model. This suggests that voltage-calcium transformation in Mi1 and Tm3 cells is different from those in T4c cells.

The time constants for the two low-pass filters were identical for the multiplicative model for T4c data fit. Thus, these two low-pass filters in the multiplicative model could also be replaced by a single low-pass filter followed by a quadratic non-linearity. An exponent of close to 2 (exact value: 2.2) was found in the parameter search for a model with a single low-pass filter followed by an exponential nonlinearity.

Differential expression of voltage-gated calcium channels in these cells could explain the different voltage to calcium transformation. Voltage-gated calcium channels mediate depolarization-induced calcium influx that drives the release of neurotransmitters. The *α*1-subunit of the voltagegated calcium channels forms the ion-conducting pore, which makes it distinct from other calcium channels. Three families of genes encode *α*1 subunits. *Drosophila* genome has one *α*1 subunit gene in each family: *α*1*D* (*Ca*_*v*_1), cac (*Ca*_*v*_2), and *α*1*T* (*Ca*_*v*_3) (Littleton & Ganetzky 2000; King 2007). In *Drosophila* antennal lobe projection neurons, cac (*Ca*_*v*_2) type and *α*1*T* (*Ca*_*v*_3) type voltage-gated calcium channels are involved in sustained and transient calcium currents, respectively (Gu *et al*. 2009; Iniguez *et al*. 2013). According to a RNA-sequencing study (Davis *et al*. 2020), *α*1*T* (*Ca*_*v*_3) mRNA have higher expression in Mi1 (2050.16 Transcripts per Million (TPM)) compared to T4 (686.68 TPM) and Tm3 (336.45 TPM). While cac (*Ca*_*v*_2) mRNA have higher expression in T4 (1298.53 TPM) compared to Mi1 (986.25 TPM) and Tm3 (817.61 TPM). Different expression of voltage-gated calcium channels could cause different voltage to calcium transformations in non-direction selective and direction-selective cells. In addition to dendritic integration of postsynaptic voltages, the specific voltage-to-calcium transformation described in this study provides an important processing step that enhances direction selectivity in the output signal of motion-sensing neurons of the fly.

## Materials and Methods

### Flies

Flies (*Drosophila melanogaster*) were raised at 25°*C* and 60% humidity on a 12 hour light/12 hour dark cycle on standard cornmeal agar medium. For calcium imaging experiments, genetically-encoded calcium indicator GCaMP6f (Chen *et al*. 2013) was expressed in T4 neurons with axon terminals predominantly in layer 3 of the lobula plate. Similarly for voltage imaging experiments, genetically-encoded voltage indicator Arclight (Jin *et al*. 2012) was expressed in T4 layer 3 neurons. The flies genotype were as follows :

1. T4c>GCaMP6f : w+ ; VT15785-Gal4AD / UAS-GCaMP6f; VT50384-Gal4DBD / UAS-GCaMP6f
2. T4c>Arclight : w+ ; VT15785-Gal4AD / UAS-Arclight; VT50384-Gal4DBD / +

For Mi1 and Tm3 experiments, the flies genotype were as follows :

1. Mi1>GCaMP6f : w+ ; R19F01-Gal4AD / UAS-GCaMP6f; R71D01-Gal4DBD / UAS-GCaMP6f
2. Mi1>Arclight : w+ ; R19F01-Gal4AD / UAS-Arclight; R71D01-Gal4DBD / +
3. Tm3>GCaMP6f : w+ ; R13E12-Gal4AD / UAS-GCaMP6f; R59C10-Gal4DBD / UAS-GCaMP6f
4. Tm3>Arclight : w+ ; R13E12-Gal4AD / UAS-Arclight; R59C10-Gal4DBD / +

### Calcium & voltage imaging

For imaging experiments, fly surgeries were performed as previously described (Maisak *et al*. 2013). Briefly, flies were anaesthetized with CO_2_ or on ice, fixed with their backs, legs and wings to a Plexiglas holder with back of the head exposed to a recording chamber filled with fly external solution. The cuticula at the back of the head on one side of the brain was cut away with a fine hypodermic needle and removed together with air sacks covering the underlying optic lobe. The neuronal activity was then measured from the optic lobe with a custom-built 2-photon microscope as previously described (Maisak *et al*. 2013). Images were acquired at 64 × 64 pixels resolution and frame rate 13 Hz with the Scanimage software in Matlab (Pologruto *et al*. 2003).

### Visual stimulation

For the study of visual responses of T4c cells, visual stimuli were presented on a custom-built projector-based arena as described in (Arenz *et al*. 2017). In brief : Two micro-projectors (TI DLP Lightcrafter 3000) were used to project stimuli onto the back of an opaque cylindrical screen covering 180° in azimuth and 105° in elevation of the fly’s visual field. To increase the refresh rate from 60 Hz to 180 Hz (at 8 bit color depth), projectors were programmed to use only green LED (OSRAM L CG H9RN) which emits light between 500 nm to 600 nm wavelength. Two long-pass filters (Thorlabs FEL0550 and FGL550) were placed in front of each projector to restrict the stimulus light to wavelengths above 550 nm. This prevents overlap between fluorescence signal and arena light spectra. To allow only fluorescence emission spectrum to be detected, a band-pass filter (Brightline 520/35) was placed in-front of the photomultiplier. Stimuli were rendered using custom written software in Python 2.7.

#### Stimuli

Stimuli were presented with 3-5 repetitions per experiment in a randomized fashion. To measure the directional and speed tuning, square-wave gratings with a spatial wavelength of 30° spanning the full extent of the stimulus arena were used. The gratings were moved in 12 different directions from 0° − 360° at 4 different speeds (15°*s*^−1^, 30°*s*^−1^, 60°*s*^−1^, 120°*s*^−1^). Similarly, to measure direction and contrast tuning, square-wave gratings with a spatial wavelength of 30° spanning the full extent of the stimulus arena were used. The gratings moved at a speed of 30°*s*^−1^ in 12 different directions at 4 different contrasts (10%, 20%, 50%, 100%). Edge responses were measured using ON edge i.e. bright edge moving on a dark background with full contrast. The ON edge moved in preferred direction (upward) or null direction (downward) at 4 different speeds (15°*s*^−1^, 30°*s*^−1^, 60°*s*^−1^, 120°*s*^−1^).

### Data analysis

Data analysis was performed using custom-written routines in Matlab and Python 2.7, 3.7. Images were automatically registered using horizontal and vertical translations to correct for the movement of brain. Fluorescence changes (Δ*F*/*F*) were then calculated using a standard baseline algorithm (Jia *et al*. 2011). Regions of interest (ROIs) were drawn on the average raw image manually by hand in the medulla layer M10 for signals from T4 dendrites. Averaging the fluorescence change over this ROI in space resulted in a (Δ*F*/*F*) time course. Voltage imaging with Arclight and calcium imaging with GCaMP6f and GCaMP8f were performed and analysed using same settings.

The direction selectivity was evaluated using a direction selectivity index (DSI) calculated as the difference of the peak responses to preferred and null direction, divided by the sum of the peak responses:

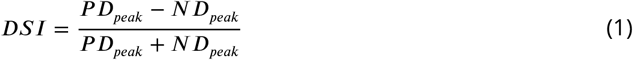

In the above measurement, only the difference in response between the two opposing directions of motion is quantified. To take into account all 12 directions of motion, we calculated the directional tuning index:

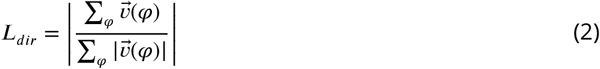

where 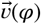 is a vector proportionally scaled with the mean response and points in the direction corresponding to the direction of motion given by the rotation angle *φ* of the stimulus (Mazurek *et al*. 2014).

### Model simulations

Custom-written Python3.7 scripts were used to simulate the models (figure 4). To calculate the optimal parameter values, we first defined an error function. For each stimulus condition (*s*_*i*_), the error was calculated as:

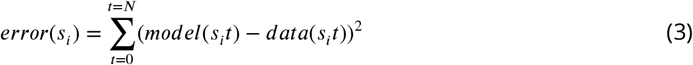

The model took as input Arclight data across all 112 different stimuli conditions. Next, we summed the error for all stimuli conditions:

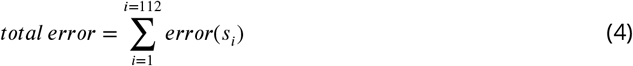

The model parameters were initialized with random values within the defined parameter bounds. Python SciPy optimize minimize function then used the L-BFGS-B (Limited Broyden Fletcher Goldfarb Shanno) algorithm to find the parameter values corresponding to the minimum total error. A total of 300 runs were performed, and the parameter values that corresponded to the run with the lowest error were used to produce the final output signals. To compare the model performances, we calculated the model error as:

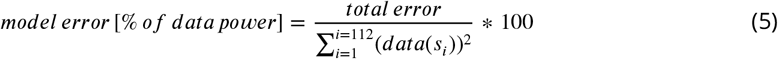

